# Loss of calbindin and rise in pT217-tau in aging monkey prefrontal cortical dendrites

**DOI:** 10.64898/2026.08.18.745599

**Authors:** Isabella J. Perone, Dinara Bolat, Zhiling Gu, Caroline Zeiss, Eliza Bliss-Moreau, Alvaro Duque, Jon Arellano, Yize Zhao, Dibyadeep Datta, Amy FT Arnsten

## Abstract

**INTRODUCTION:** Tau pathology in Alzheimer’s disease preferentially afflicts excitatory neurons in the limbic and association cortices that utilize high levels of calcium signaling to perform cognitive operations. This includes the layer III pyramidal cells in the dorsolateral prefrontal cortex (dlPFC) that subserve higher cognition, which express the calcium-binding protein, calbindin, when young and healthy, but lose calbindin and develop tangles and degenerate in Alzheimer’s disease (AD). These data suggest that loss of calbindin may be associated with the emergence of tau pathology. However, the relationship between calbindin and early-stage, soluble tau pathology is challenging to study in human brains, as soluble pTau dephosphorylates within 15min postmortem. In contrast, the relationship between calbindin and soluble pT217-tau expression can be studied in aging macaques with naturally-occurring tau pathology, where perfusion fixation is possible to capture phosphorylation state in situ.

**METHODS:** The current study used multiple-label-immunofluorescence to label MAP2-positive dlPFC layer III pyramidal cells for calbindin and pT217-tau in macaque brains across the adult age span (8-34.5yrs). The study employed a semi-automated CellProfiler workflow to identify labeled pyramidal cell dendrites the cellular compartment where tau pathology begins in AD.

**RESULTS:** Calbindin expression decreased with age, while pT217Tau increased with age. Specifically, the ratio of calbindin/pT217-tau within a dendrite decreased with age, and was especially prominent in the aged macaques with long-term inflammatory disorders.

**DISCUSSION:** These data suggest that the loss of calbindin in dendrites with advancing age, and especially with inflammation, contributes to the rise of tau pathology and the risk of AD.

## BACKGROUND

Alzheimer’s disease (AD) selectively impacts higher cortical circuits, with tau pathology and neurodegeneration preferentially afflicting glutamatergic neurons in limbic and association cortices^1–2^. Neuropathological studies have shown that AD-related neurofibrillary changes in cortex progress in a stereotyped anatomical sequence, beginning in entorhinal regions, extending through hippocampal and limbic structures and association cortices, and ultimately reaching primary sensori-motor cortices^1–2^]. Early tau pathology also shows subcellular specificity, with abnormal tau first appearing in distal dendritic compartments and proceeding toward the soma^3^. Thus, understanding the etiology of AD requires identifying the molecular events that render select higher cortical circuits and their dendritic compartments vulnerable during aging, before the emergence of end-stage tangles and overt neuronal loss.

Aging rhesus macaques provide a unique window into early-stage AD-like pathology in higher cortical circuits^4–6^. Unlike rodents, rhesus macaques possess extensive association cortices, are ApoE-ε4 homozygotes, and naturally develop age-related tau and amyloid pathology, synapse loss, and cognitive deficits^4–6^. Importantly, perfusion-fixed macaque tissue permits visualization of early, soluble phosphorylated tau species that are rapidly dephosphorylated in human postmortem tissue^4,7^. This is especially important for tau phosphorylated at threonine 217 (pT217Tau), a soluble p-tau species that has emerged as a powerful fluid biomarker of incipient AD pathology^8^. Nanoscale imaging of aged macaque entorhinal cortex and dorsolateral prefrontal cortex (dlPFC) has shown that early-stage, soluble pT217Tau aggregates on microtubules within dendrites, and can traffic between excitatory neurons at glutamatergic synapses^9^. These findings suggest that soluble pT217Tau may seed pathology across vulnerable cortical circuits, and can gain access to CSF and plasma as a fluid biomarker^9^.

A growing body of work suggests that inflammation with advancing age drives calcium dysregulation, which in turn promotes early tau pathology, APP processing, and Aβ generation^5,10,11^. The dlPFC expresses magnified calcium signaling needed to maintain neuronal firing for memory and cognitive operations^5^. This must be tightly regulated, e.g. by the calcium binding protein calbindin, to prevent toxic actions^5^. However, regulation is lost with age and/or inflammation^5^. Very high levels of cytosolic calcium activate calpain-2, promoting tau hyperphosphorylation through downstream kinases such as GSK3β and cdk5, while also contributing to APP cleavage, Aβ generation, and autophagic degeneration^4,5^. These processes may form vicious cycles in which p-tau, Aβ, inflammation, and excessive calcium signaling drive one another across the aging brain^4,5^. Consistent with this model, restoring calcium regulation in aged macaque cortex reduces pT217Tau levels, suggesting that calcium dysregulation is a modifiable driver of early-stage tau pathology^12^.

Calbindin-D28K is central to this framework, because it helps regulate cytosolic calcium in neurons with high calcium demands^13^. However, calbindin is lost from many neurons with advancing age^13–14^, and especially with inflammation^14^. In macaques, calbindin is expressed in the layer III dlPFC pyramidal cells that generate working memory^16^, but is lost with age^14^. In human dlPFC, calbindin-labeled layer III pyramidal neurons are especially afflicted in AD, with their loss correlating with neurofibrillary tangle burden^17^. In contrast, superficial interneuron populations with higher calbindin expression are relatively spared in both macaques and humans^14,17^. The loss of calbindin from layer III pyramidal cells with advancing age, while remaining relatively preserved in interneurons, may help explain the selective vulnerability of these neurons to tau pathology^5,14^.

The current study examined the relationship between pT217Tau and calbindin in adult rhesus macaque dlPFC layer III pyramidal cell dendrites across the age span. Our data suggest an age-related overall increase in pT217Tau signal, a decrease in calbindin, and a reduction in the calbindin/pT217Tau ratio within dlPFC dendrites. These findings support a model in which aging shifts vulnerable layer III pyramidal neurons away from calcium-buffering and toward a pathological state marked by rising tau phosphorylation and declining calcium-regulation. By focusing on dendrites, the present work examines a subcellular compartment where tau pathology and synaptic dysfunction may emerge early in disease^3^.

## MATERIAL AND METHODS

### Animals and Tissue

Ten adult rhesus macaques (*Macaca mulatta*) spanning the adult age range were included in this study (Supplemental Table 1). Animals ranged from 8.02 to 34.49 years of age, encompassing young adult, middle-aged, and aged cohorts. This range corresponds roughly to the third through seventh decades of human life, a period that encompasses normal aging through the emergence of preclinical AD-like pathology in cortical circuits^21^.

Tissue was acquired at three sites. Eight animals were processed at the primary acquisition facility (Site 3, reference cohort); one was processed at Site 1 (8.02 years) and one at Site 2 (11.28 years). Because multi-site processing introduces systematic immunofluorescence differences independent of age, site of acquisition was treated as a fixed covariate in all statistical models (Supplemental Methods S1).

### Tissue processing

The animals used in this study and were maintained and euthanized in accordance with the guidelines of the US Department of Agriculture, the Yale University Institutional Animal Care and Use Committee (IACUC), and in accordance with the Weatherall report. As described previously^6,18–20^, primates were deeply anesthetized prior to transcardial perfusion of 100 mM phosphate-buffered saline, followed by 4% paraformaldehyde/0.05% glutaraldehyde in 100 mM phosphate-buffered saline. After perfusion, a craniotomy was performed, and the entire brain was removed and blocked. The brains were sectioned coronally at 60 μm on a vibratome (Leica) across the entire rostrocaudal extent of the dlPFC (Walker’s area 46). The sections were cryoprotected through increasing concentrations of sucrose solution (10%, 20%, and 30% each for overnight), cooled rapidly using liquid nitrogen, and stored at −80°C. Non-specific reactivity was suppressed with 10% normal goat serum (NGS) and 5% bovine serum albumin (BSA), and antibody penetration was enhanced with 0.3% Triton X-100 in 50 mM Tris-buffered saline (TBS).

### Immunoreagents

Previously characterized primary antibodies were used, and complexed with species-specific goat secondary antibodies. For pT217Tau ^9^: rabbit anti-pT217Tau at 1:100 (cat# AS-54968, Anaspec, RRID:AB_2173656). The immunogen used keyhole limpet hemocyanin conjugated with synthetic peptides corresponding to human tau at phosphorylated threonine 217. The antiserum was produced against a chemically synthesized phosphopeptide derived from the region of human tau that contains threonine 217. For CB^14^: mouse anti-calbindin D-28k at1:500 [cat# 300, Swant Antibodies, RRID:AB_10000347]. The specificity and selectivity of the primary antibodies have been characterized using immunoblot, immunohistochemistry, immunofluorescence, and enzyme-linked immunosorbent assay approaches in various tissue types in multiple species^1,6,22, 23^. Normal sera and immunoglobulin G (IgG)-free BSA were purchased from Jackson ImmunoResearch.

### Immunofluorescence

Free-floating dlPFC sections were processed for triple-label immunofluorescence as described previously^19^. Antigen retrieval was performed with 10 mM sodium citrate buffer (pH 6.0) in a hot water bath (∼75°C) for 25 min, followed by 30 min cooling at room temperature. Sections were transferred to 1× TBS for 10 min, then incubated in 0.5% sodium borohydride in 0.01 M TBS to reduce background autofluorescence. Sections were blocked for 1 h at room temperature in 1× TBS containing 5% bovine serum albumin, 2% Triton X-100, and 10% normal goat serum. Primary antibodies — anti-MAP2 [dilution 1:1000; ab5392, Abcam], anti-calbindin D-28k at 1:500 [cat# 300, Swant Antibodies], and anti-pT217Tau at 1:100 [cat# AS-54968, Anaspec]. — were applied for 48 h at 4°C. Sections were then incubated overnight at 4 °C with species-appropriate, cross-adsorbed goat secondary antibodies (all 1:1,000): Alexa Fluor 488 goat anti-mouse IgG (H+L) (Cat# A-11001; RRID:AB_2534069), Alexa Fluor 594 goat anti-rabbit IgG (H+L) (Cat# A-11012; RRID:AB_2534079), and Alexa Fluor 633 goat anti-chicken IgY (H+L) (Cat# A-21103; RRID:AB_2535756; Invitrogen). To suppress lipofuscin autofluorescence, sections were treated with 70% ethanol containing 0.3% Sudan Black B (MP Biomedicals, Cat# 4197-25-5). Nuclei were counterstained with Hoechst 33342 (1:10,000; Thermo Fisher, Cat# H3570) and sections were mounted using ProLong Gold Antifade Mountant (Invitrogen, Cat# P36930). Confocal images were acquired on a Zeiss LSM 880 Airyscan Confocal Microscope using the C-Apochromat 40×/1.2 W Korr FCS M27 water objective (Zeiss). Sequential scanning with matched emission filter bandwidths was used to avoid spectral overlap between fluorophores. Images were merged using Fiji. Approximately 10–15 images were collected per cortical layer per section, capturing approximately 15 pyramidal cells per image at 40× magnification. In order to capture a consistent number of images, 2-3 sections per animal were analyzed.

### Semi-Automated Image Analysis: CellProfiler Pipeline Overview

Quantitative analysis of calbindin and pT217Tau immunofluorescence was performed using a customized semi-automated CellProfiler pipeline (version 5.x; Broad Institute)^24^. The pipeline was designed to isolate and quantify fluorescence signals specifically within MAP2-positive dendritic compartments, ensuring that all measurements reflect dendritic protein distribution.

The pipeline proceeded through the following steps:

#### Step 1: Nuclear identification

Cell nuclei were identified from the Hoechst/DAPI channel (OrigBlue) using the IdentifyPrimaryObjects module. Objects with diameters between 40 and 400 pixels were retained. A global two-class Otsu thresholding strategy was applied to the log-transformed image with a correction factor of 1.25, threshold smoothing scale of 1.3488, and intensity bounds of 0.05–1.0. The resulting segmented objects (Nuclei) provided the nuclear mask used for soma approximation and exclusion in downstream steps (Figure 1A).

**Figure 1.**
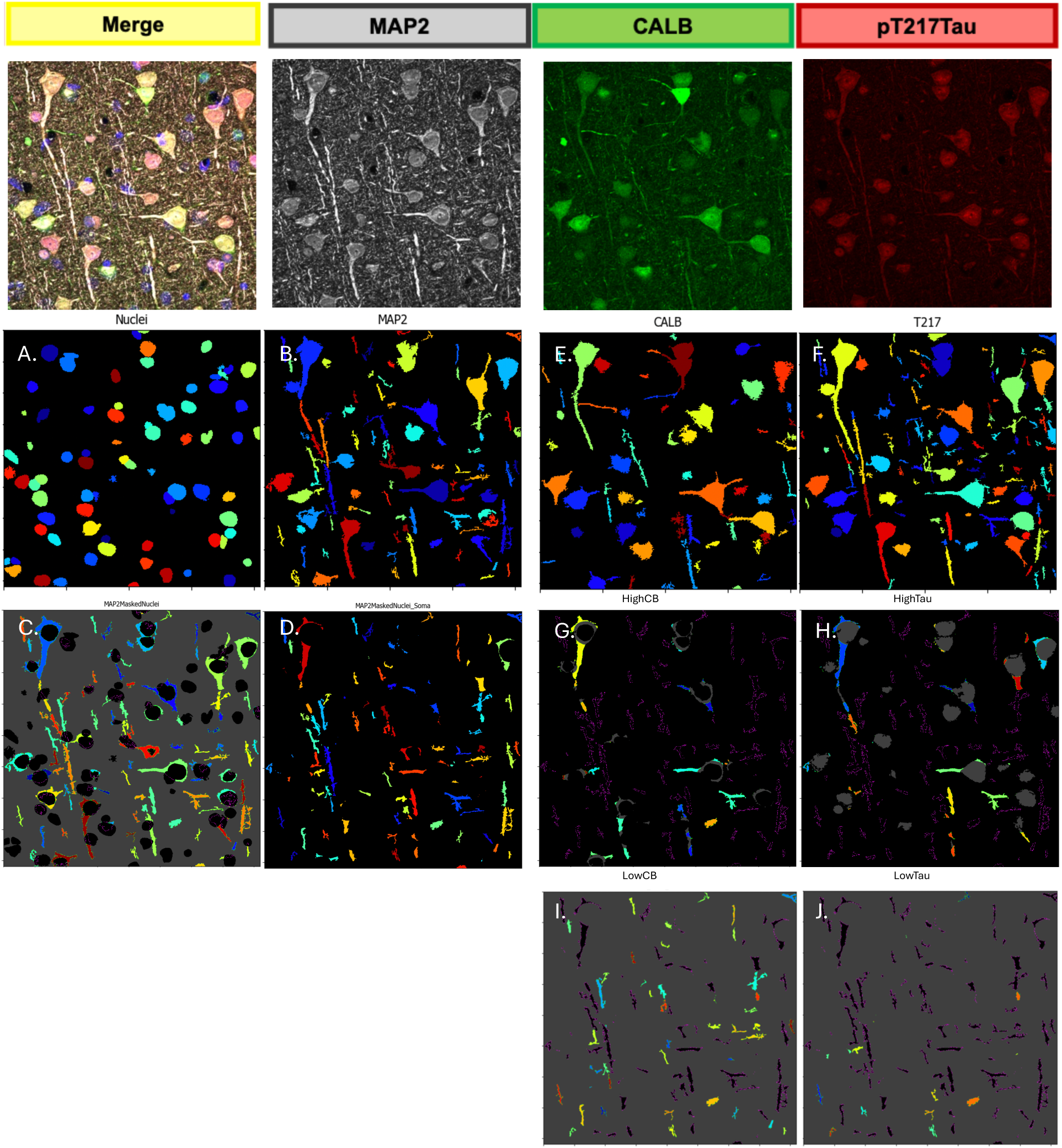
Representative frames from CellProfiler pipeline. Top panel shows the original images fed into pipeline. Panels A-D highlight nuclear labeling (A), MAP2 labelling (B), and the masks used to block the nucleus (C) analyze exclusively within dendrites (D). Panels E-J show masks selecting for calbindin (E), and pT217Tau (F) positivity respectively, as well as representative masks showing both the highest (G-H) and lowest (I-J) intensity threshold groups for both markers.

#### Step 2: MAP2-positive object detection

All MAP2-immunoreactive structures, including neuronal cell bodies and dendritic compartments, were identified from the MAP2 channel (OrigWhite) using the IdentifyPrimaryObjects module. The diameter range was set to 30–10,000 pixels to capture both dendritic segments and full cell bodies without size-based exclusion. A global three-class Otsu strategy was applied to the log-transformed image (middle class assigned to foreground; correction factor 1.5; bounds 0.25–1.0) with a large smoothing filter (10,000 px) that treated the MAP2-positive field as a continuous region (Figure 1B).

#### Step 3: Dendritic mask definition

The Nuclei objects were expanded radially by 25 pixels to approximate the neuronal soma boundary. Two complementary dendritic masks were then generated by applying inverted masks to MAP2 objects: MAP2MaskedNuclei retains MAP2-positive signal outside the nucleus (soma cytoplasm included) (Figure 1C), while the expanded mask retains only dendritic processes outside the expanded soma boundary. The expanded mask served as the primary analysis compartment for all fluorescence intensity quantification. In both masks, objects partially overlapping the mask boundary were retained with a minimum overlap threshold of 0.5 (Figure 1D).

#### Step 4: Calbindin and pT217Tau segmentation and intensity stratification

Within the dendritic mask, intensity-based thresholds were independently applied to calbindin and pT217Tau channels. For calbindin (Figure 1E), a base segmentation (Minimum Cross-Entropy 3-class; correction factor 1.25; bounds 0.09–1.0) anchored the per-image reference and four bins were defined: high calbindin (≥0.301), above-average (0.213–0.301), below-average (0.125–0.213), and low (0.090–0.125). Each of the eight intensity bins was intersected with the dendritic mask to produce dendritic-compartment-restricted object sets. Both per-dendrite mean fluorescence intensity values and the proportion of dendritic area occupied by each intensity category were recorded. For pT217Tau (Figure 1F), an initial segmentation (global Otsu 3-class, log-transformed; correction factor 1.7; bounds 0.20–1.0) established the per-image reference threshold. Four mutually exclusive intensity bins were then defined relative to this reference: high pT217Tau (≥0.455), above-average (0.345–0.455), below-average (0.234–0.345), and low (0.200–0.234). Representative frames from the highest and lowest threshold groups for both calbindin and pT217Tau are represented in Figure 1G-J.

### Intensity Thresholding and Area-Occupied Quantification

To quantify the spatial footprint of each marker within the dendritic compartment, fluorescence intensity within the dendritic mask was binned into four mutually exclusive intensity categories: Low, Below Average, Above Average, and High, using automated, consistently applied thresholds. Threshold bounds were calibrated empirically against a reference subset of images prior to batch processing: for pT217Tau, the high-intensity lower bound (≥0.455) was set to correspond to the mean Otsu threshold in aged animals plus approximately one standard deviation; for calbindin, the high-intensity lower bound (≥0.301) was calibrated against mean calbindin signal in a young reference group. All thresholds were finalized prior to batch analysis and held constant across the full cohort. For each category, the area (in pixels) occupied within the dendritic mask was summed per animal across all images and expressed as a percentage of total categorized dendritic area.

### Statistical Analysis

Two complementary statistical frameworks were applied, both analyzing the same dendritic compartment but capturing distinct dimensions of marker expression.

#### Dendritic intensity analysis

Per-image correlations between animal age and mean dendritic calbindin or pT217Tau intensity were computed to characterize trends. At the per-dendritic object level, linear mixed-effects models (MEMs) were fitted in R using the lme4 and lmerTest packages with the model structure: outcome ∼ age + site + (1|animal/image). The nested random-intercept structure accounts for the non-independence of dendritic objects within images and images within animals. Site was included as a fixed additive covariate. In total, 17,910 dendritic objects from 263 images across 10 animals were analyzed. FDR correction was applied across models within each marker family.

#### Dendritic area-occupied analysis

For area-occupied outcomes, the unit of analysis was per animal (n = 10). Site-adjusted ordinary least-squares (OLS) regression was applied: outcome ∼ age + site, with Site 3 as the reference. Three CB/pT217Tau ratio outcomes were examined: (1) High CB / High pT217Tau, (2) (High + Above Average CB) / High pT217Tau, and (3) (High + Above Average CB) / (High + Above Average pT217Tau). FDR correction was applied within the ratio family and separately within the CB and pT217Tau individual category families. All analyses were conducted in R. Statistical significance was set at α = 0.05 after FDR correction. The statistical approach was developed in consultation with Dr. Yize Zhao (Yale School of Public Health). Details of the site correction procedure are provided in Supplemental Methods S1.

## Results

### Representative Immunofluorescence across the Adult Age Span

Triple-label immunofluorescence for MAP2, calbindin (CB), and pT217Tau was examined in dlPFC layer III sections from ten rhesus macaques spanning the adult age range (ages 8.02–34.49yrs). Representative fields from each animal are shown in Figures 2–4, organized by age group: young/middle-aged adults (8–11yrs; Figure 2), early aged adults (18–19yrs; Figure 3), and older aged adults (23–34yrs; Figure 4). As expected from previous research, young/middle-aged animals displayed strong, uniform CB immunoreactivity with especially high CB immunoreactivity in GABA interneurons and lower intensity of CB immunoreactivity of MAP2-positive pyramidal cells in dlPFC layer III. These animals also showed a subtle pT217Tau immunoreactivity in MAP2-positive dendritic compartments (Fig. 2). The early aged macaques exhibited an intermediate pattern of CB and pT217Tau immunoreactivity, with notable heterogeneity across fields (Fig 3). Examples of dendrites from the 19yrs macaque are shown at higher magnification in Figure 5, emphasizing the divergence emerging with advancing age, including the appearance of pT217Tau in many dendrites devoid of CB or with subtle CB immunoreactivity. The oldest aged animals (23-34yrs) generally showed reduced CB expression and increased pT217Tau immunoreactivity in MAP2-positive dendritic compartments (Fig. 4), with this pattern particularly evident in the two animals with chronic inflammatory conditions (23, 31yrs) (Supplementary Table 1). Intriguingly, in the oldest animal (34yrs), all labeling was less intense, as analyzed below. The heterogeneity in age-related changes is consistent with can be observed in aging humans and rodents and can help to reveal the conditions associated with vulnerability vs. resilience to emerging tau pathology^25^.

**Figure 2.**
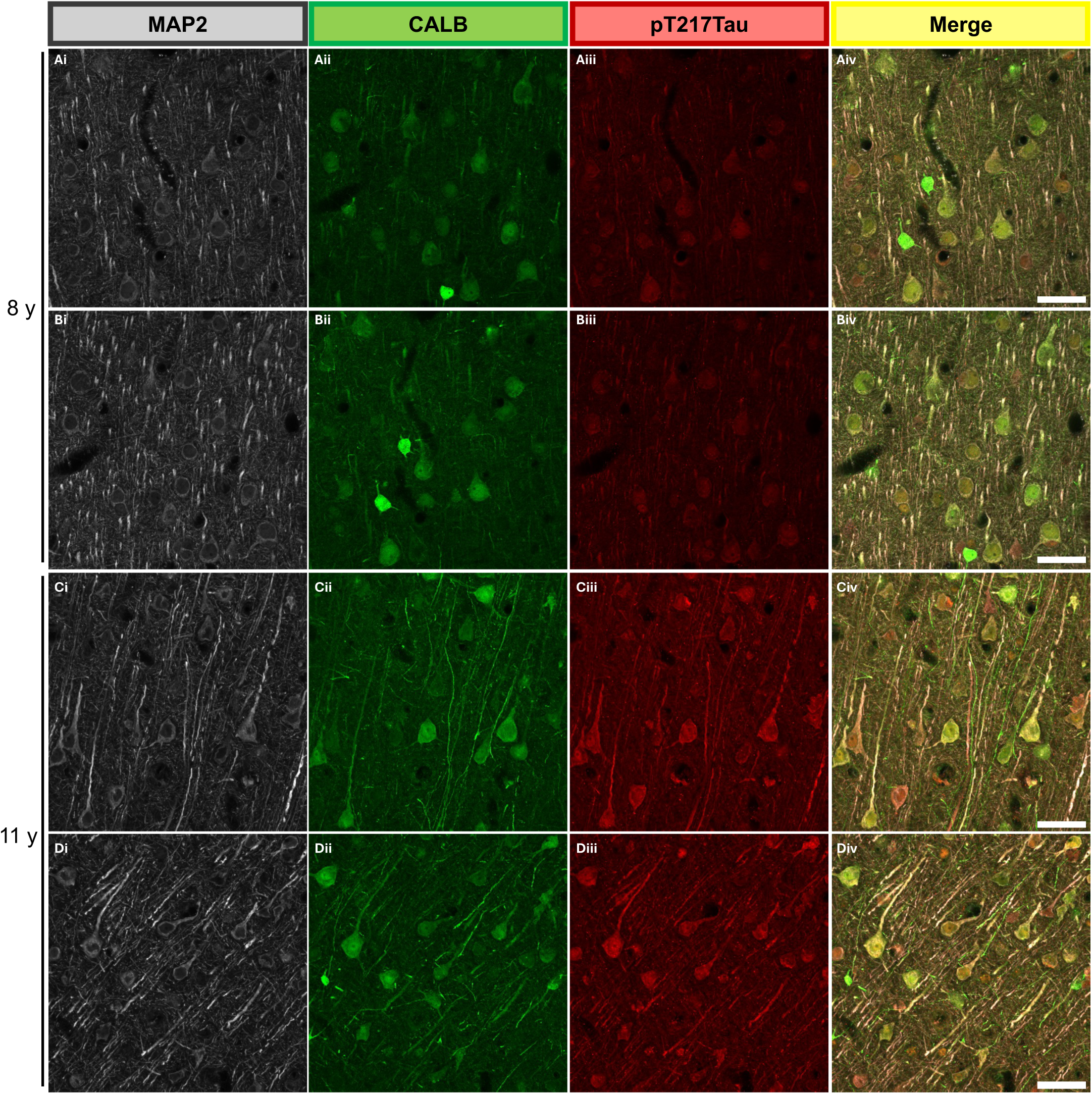
Representative triple-label immunofluorescence in dlPFC layer III MAP2-positive dendrites — middle aged adult macaques (8y, 11y). Columns show MAP2 (grey), calbindin (CB; green), pT217Tau (red), and merged composite. Each pair of rows shows two image fields from one animal. Scale bars = 40um.

**Figure 3.**
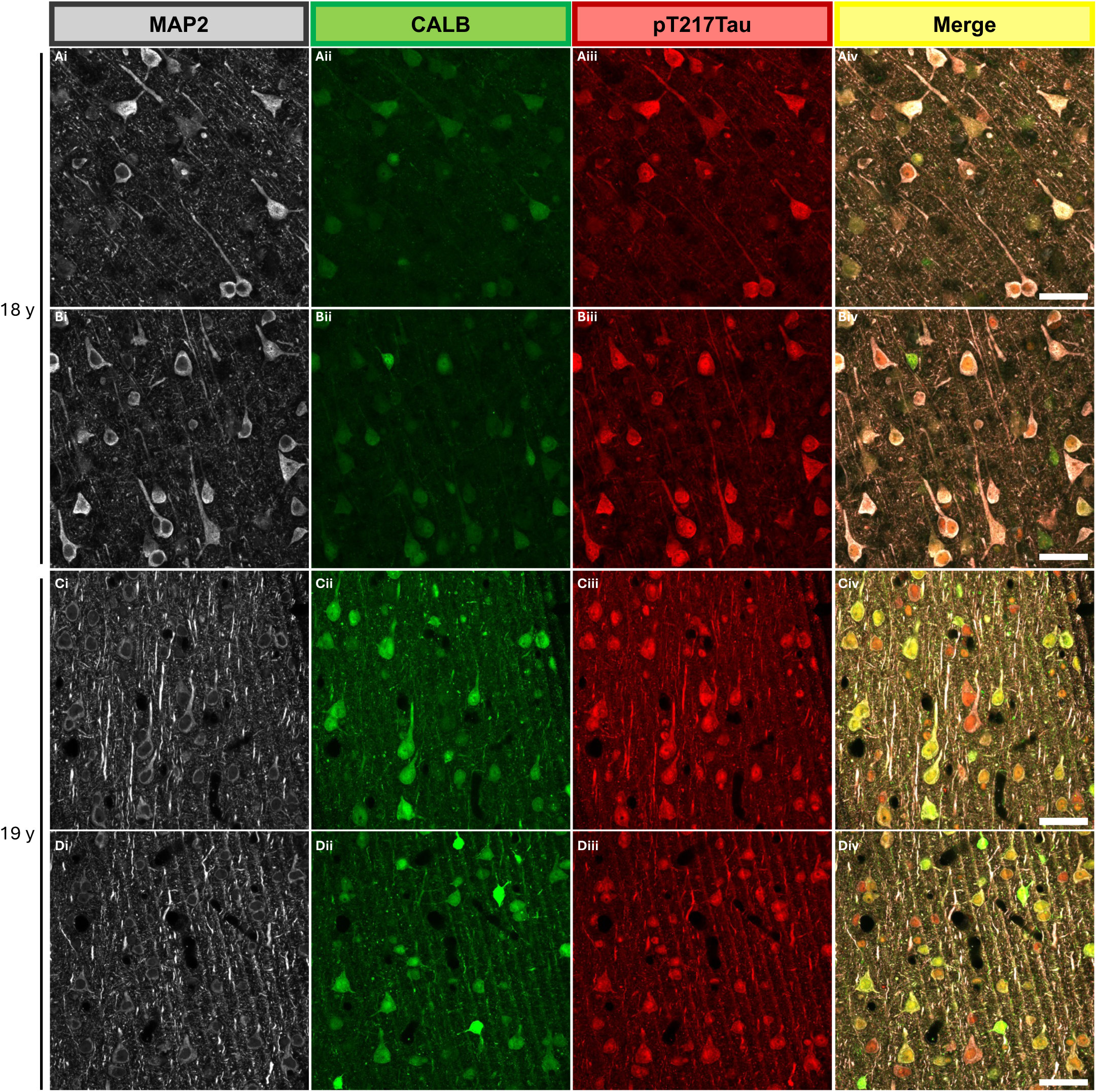
Representative triple-label immunofluorescence — early aged adult macaques (18y, 19y). Columns show MAP2 (grey), calbindin (CB; green), pT217Tau (red), and merged composite. Each pair of rows shows two image fields from one animal. Relative to middle aged adult animals, an intermediate pattern of calbindin and pT217Tau immunoreactivity is observed, with some heterogeneity across fields. Scale bars = 40 um

**Figure 4.**
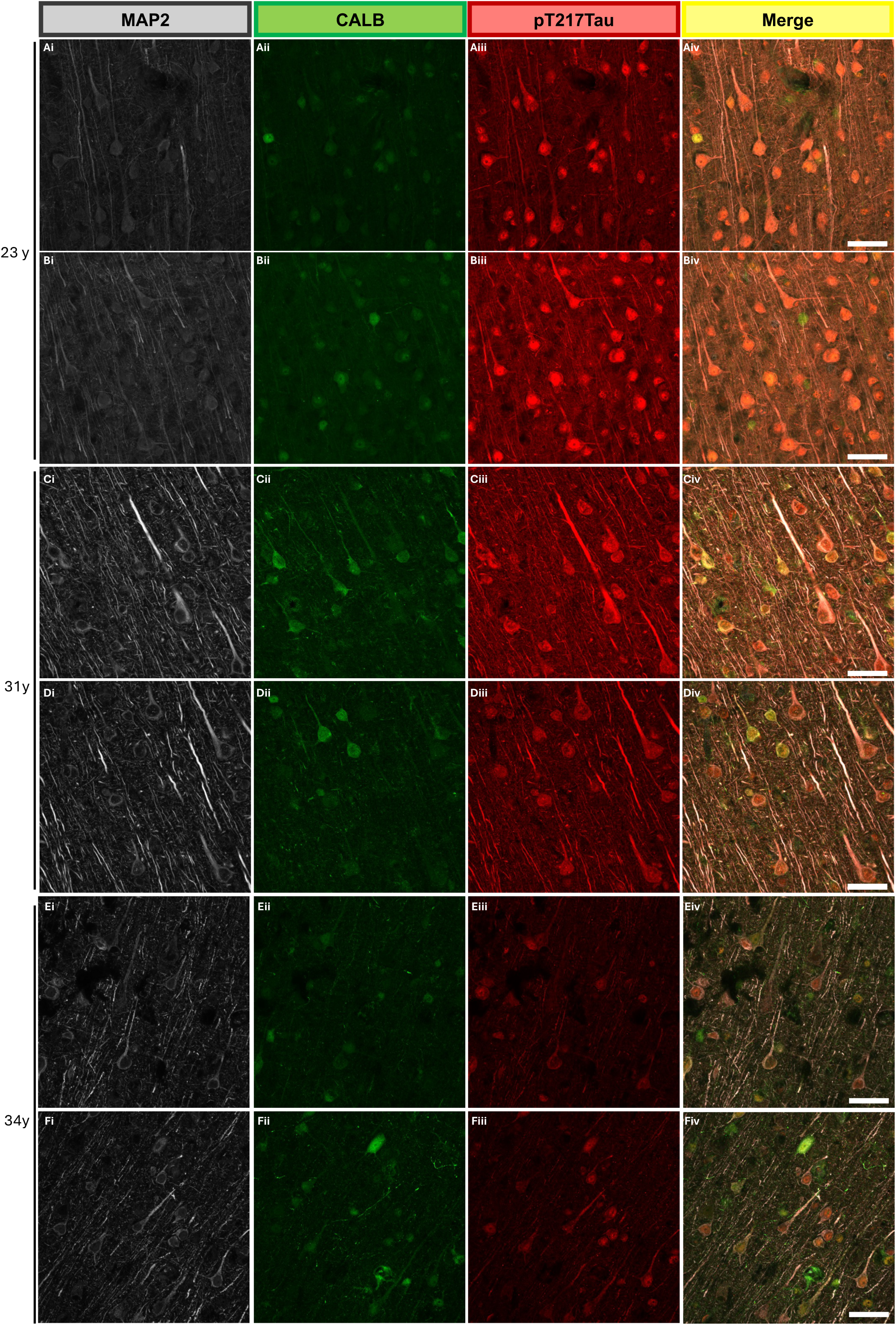
Representative triple-label immunofluorescence — older aged adult macaques (23y, 31y and 34y). Columns show MAP2 (grey), calbindin (CB; green), pT217Tau (red), and merged composite. Each pair of rows shows two image fields from one animal. Animals aged 23y and 31y with chronic inflammatory disorders show markedly reduced calbindin immunoreactivity and increased pT217Tau immunoreactivity within MAP2-positive dendritic compartments, consistent with the age-related calcium dysregulation driving tau phosphorylation, whereas animal aged 34y shows reduced calbindin immunoreactivity and limited pT217Tau signal within MAP2-positive dendritic compartments, in contrast with the expected age-related trend. Scale bars = 40um.

**Figure 5.**
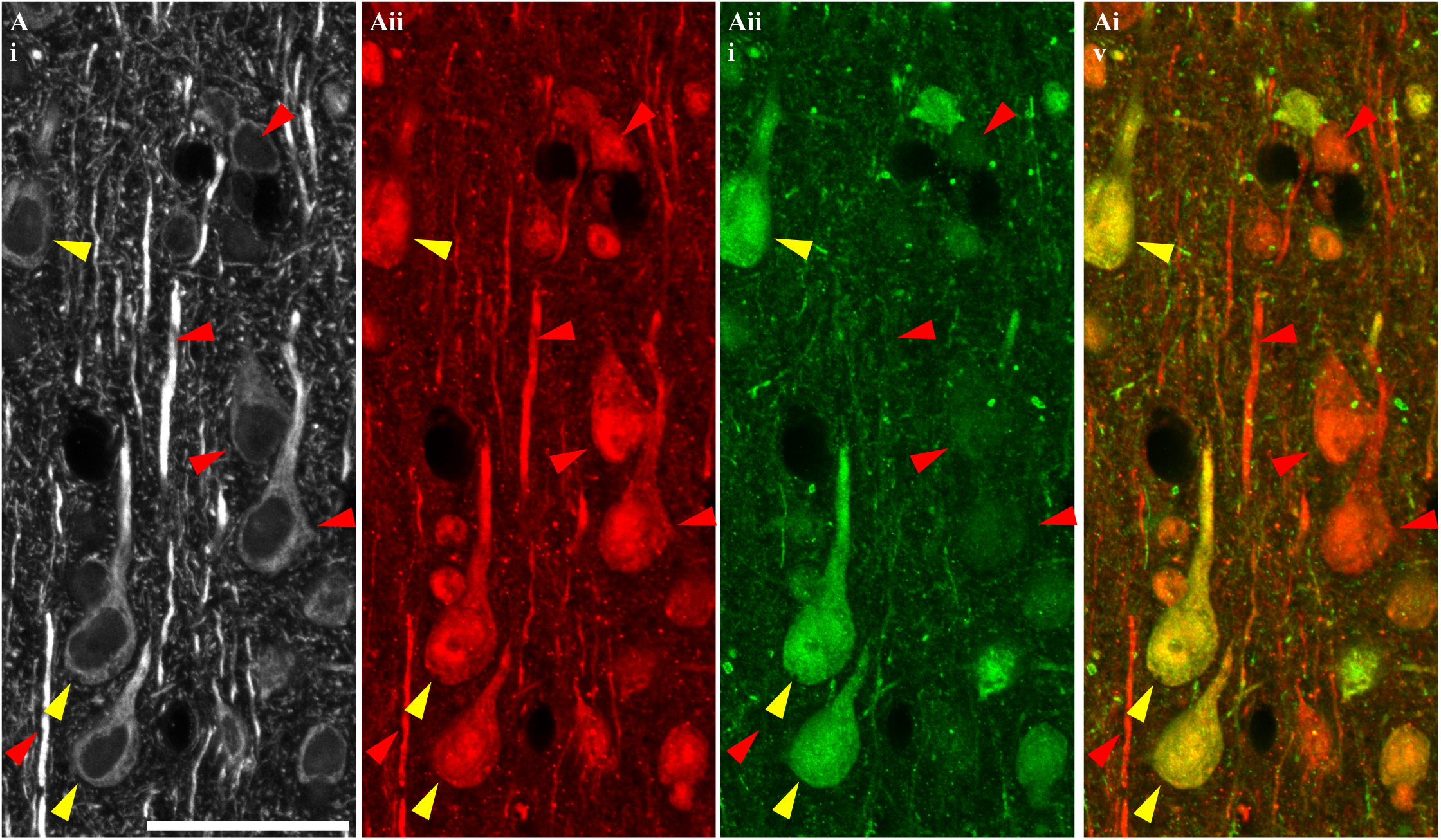
Higher-magnification representative fields across from an early aged animal (.19yrs). Shows calbindin (CB; green), pT217Tau (red), and merged composite (yellow). Red triangles indicate cells and dendrites positive for pT217Tau that are devoid of CB and, green triangles indicate cells and dendrites that are CB positive and devoid of tau. The range of co-expression patterns across and within animals illustrates the degree of heterogeneity in pT217Tau accumulation and calbindin loss captured by across the age span. Scale bars = 40um

### Calbindin-D28K and pT217Tau Dendritic Area Occupancy Across the Adult Lifespan

To characterize how the balance between CB and pT217Tau immunoreactivity shifts within layer III pyramidal cell dendritic compartments of the aging dlPFC, an area-occupied analysis was performed across the 10 rhesus macaques. The soma and perisomatic space were masked out of all images prior to analysis (Figure 1), restricting measurements exclusively to the dendritic compartment — the subcellular site where tau pathology begins in AD^3^, and where soluble pT217Tau has been shown to first accumulate on microtubules in aging macaque cortex^9^. For each marker, the proportion of dendritic compartment area occupied by four intensity categories — Low, Below-Average, Above-Average, and High — was quantified per animal and modeled as a function of age, with acquisition site included as a covariate to account for tissue processing differences across three sites (OLS: outcome ∼ age + site; n = 10).

### Calbindin-D28K intensity categories

Across the lifespan, there was a consistent trend toward declining high-intensity CB occupancy with advancing age. The proportion of dendritic compartment area in the High CB category decreased by 0.121 percent on average for every year of a given animals age (SE = 0.143, p = 0.432; Figure 6A-D), and the cumulative High + Above-Average CB fraction declined at 0.538 percent per year (SE = 0.411, p = 0.238; FDR-p = 0.534). Below-average and low-intensity CB fractions showed no consistent directional trend (all p > 0.32). None of the individual CB category comparisons survived FDR correction (all FDR-p > 0.53), consistent with the modest effect sizes and the scarcity of aged macaque tissue that constrains cohort size in this model^26^.

**Figure 6.**
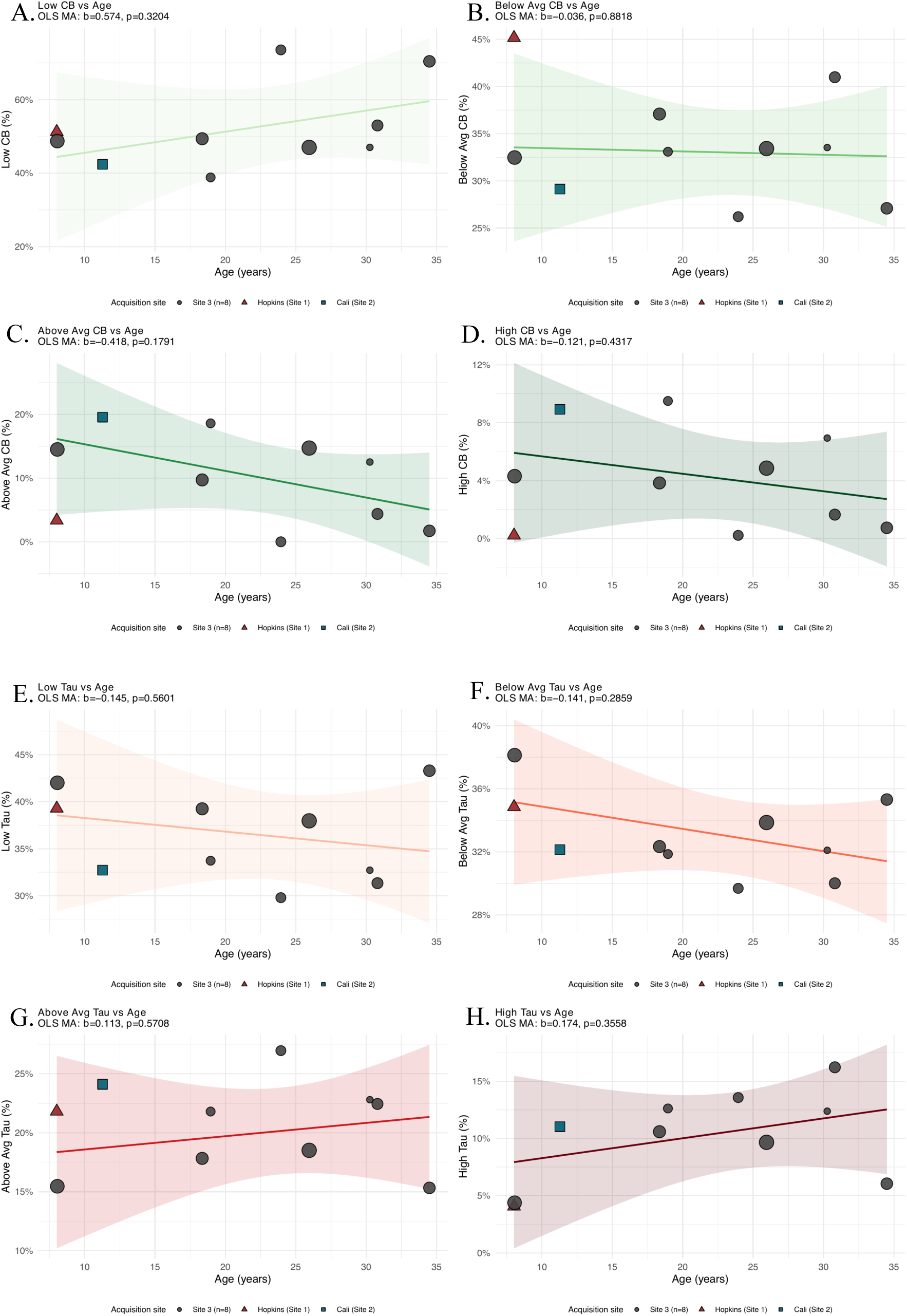
Calbindin-D28K intensity category area occupancy vs. age across 10 rhesus macaques. Each panel shows the percentage of dendritic compartment area in a given CB (Figure 6A-D) or pT217Tau (Figure 6E-H) intensity category (Low, Below-Average, Above-Average, High) as a function of age (years). Points are shaped and colored by acquisition site. Line = site-adjusted OLS fit (Site 3 reference) ± 95% CI.

### pT217Tau intensity categories

In contrast to CB, high-intensity pT217Tau area occupancy trended upward with advancing age. The proportion of dendritic compartment area in the High pT217Tau category increased at a rate of 0.174 percent (SE = 0.174, p = 0.356; Figure 6E-H), and the cumulative High + Above-Average pT217Tau fraction showed a similar positive trend (β = +0.286 %/year, SE = 0.347, p = 0.440). Low- and below-average-intensity pT217Tau fractions showed minimal directional change (all p > 0.29). As with CB, no individual pT217Tau intensity category reached FDR-corrected significance across the 10-animal cohort (all FDR-p > 0.57), due to the limited statistical power conferred by n = 10 and the substantial individual variability in tau burden at any given age.

It is noteworthy that the oldest animal (34yrs) had fainter labeling in all channels, and thus differed from the other old aged animals in the pattern of its tau labeling. This animal showed elevated pT217Tau in the low and below-average tau intensity categories, rather than the high intensity categories represented in other old aged animals. However, this variability between animals in CB and pT217Tau expression was decreased when the images were analyzed for the relationship between CB and pT217Tau within individual dendrites.

### CB/pT217Tau area-occupied ratios

The ratio of CB and pT217Tau signals within the same dendritic compartment revealed a robust and consistent age-related decline. All three CB/pT217Tau area-occupied ratios declined significantly with age following correction for acquisition site, and all three survived FDR correction within the ratio outcome family (FDR-p = 0.037; Figure 7). The ratio of the percent of High-intensity CB to High-intensity pT217Tau area (SE = 0.0099, p = 0.026; FDR-p = 0.037), indicating that high-intensity CB area shifts to high-intensity pT217Tau area as a function of age. The broader ratio of High + Above-Average CB to High pT217Tau area showed the most pronounced decline (β = –0.124/year, SE = 0.036, p = 0.013; FDR-p = 0.037), and the ratio of High + Above-Average CB to High + Above-Average pT217Tau area likewise declined significantly (β = –0.029/year, SE = 0.011, p = 0.037; FDR-p = 0.037). The youngest animal in the cohort (Site 1) showed substantially lower CB/pT217Tau ratios than expected for its age relative to Site 3 animals, consistent with a likely perfusion-related processing difference (High CB / High pT217Tau site offset: β = –0.770, SE = 0.284, p = 0.035); correcting for this site effect sharpened the age estimate for ratio outcomes, as detailed in the Methods. Site 2 showed no significant offset for any ratio outcome (all p > 0.42). Together, these findings demonstrate a progressive, age-related shift in the balance of calbindin-D28K and pT217Tau immunoreactivity within dlPFC layer III pyramidal cell dendritic compartments — with high-intensity CB area declining relative to high-intensity pT217Tau area across the macaque adult lifespan — consistent with a model in which diminishing calcium-buffering capacity accompanies rising tau phosphorylation in circuits selectively vulnerable in AD.

**Figure 7.**
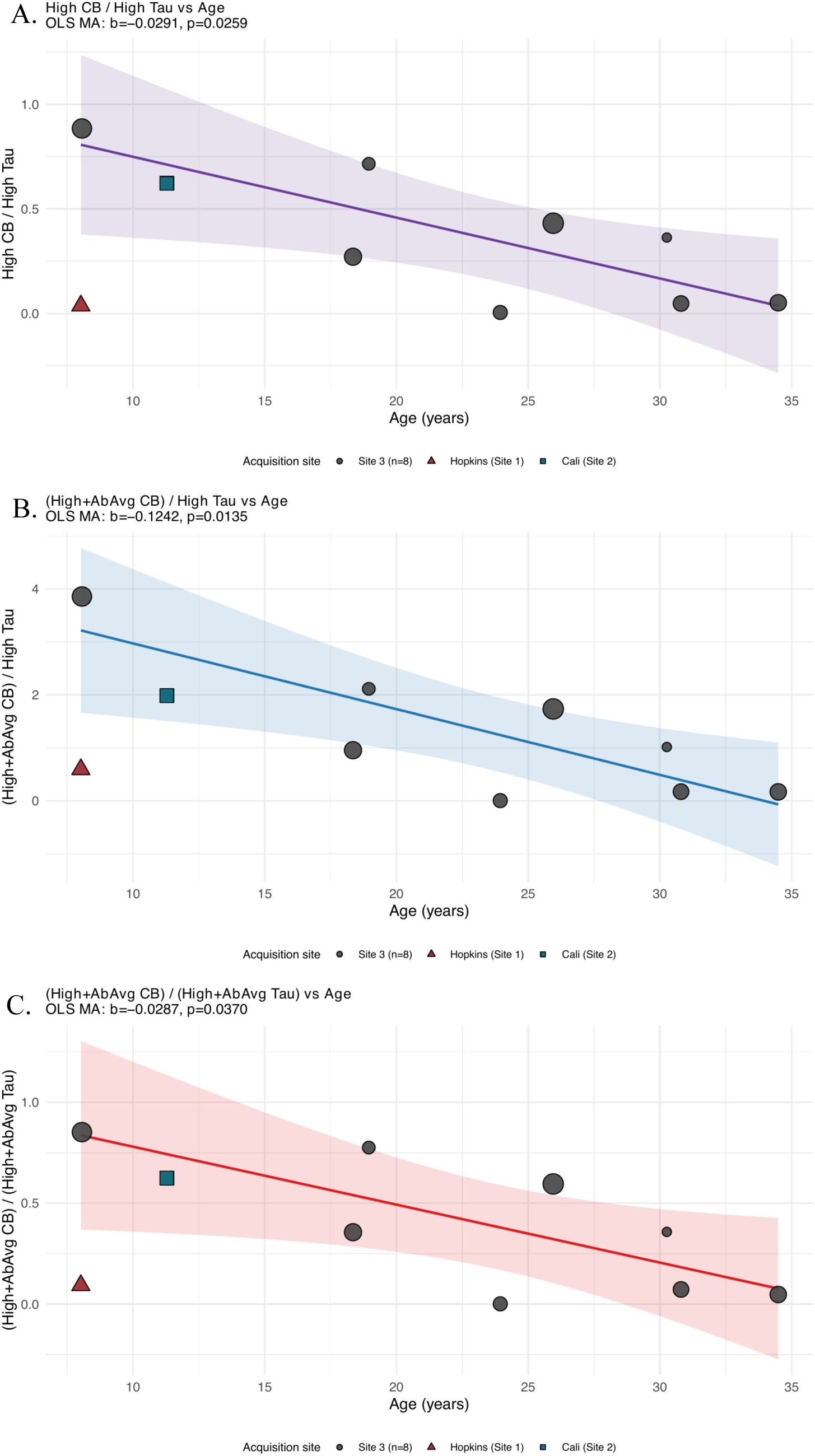
CB/pT217Tau area-occupied ratios vs. age across 10 rhesus macaques. Three ratio definitions are shown: A. High CB / High pT217Tau (top), B. (High + Above Average CB) / High pT217Tau (middle), and C. (High + Above Average CB) / (High + Above Average pT217Tau) (bottom). All three decline significantly with age after site correction (FDR-p = 0.037). Points are shaped and colored by acquisition site. Line = site-adjusted OLS fit (Site 3 reference) ± 95% CI.

**Figure 8.**
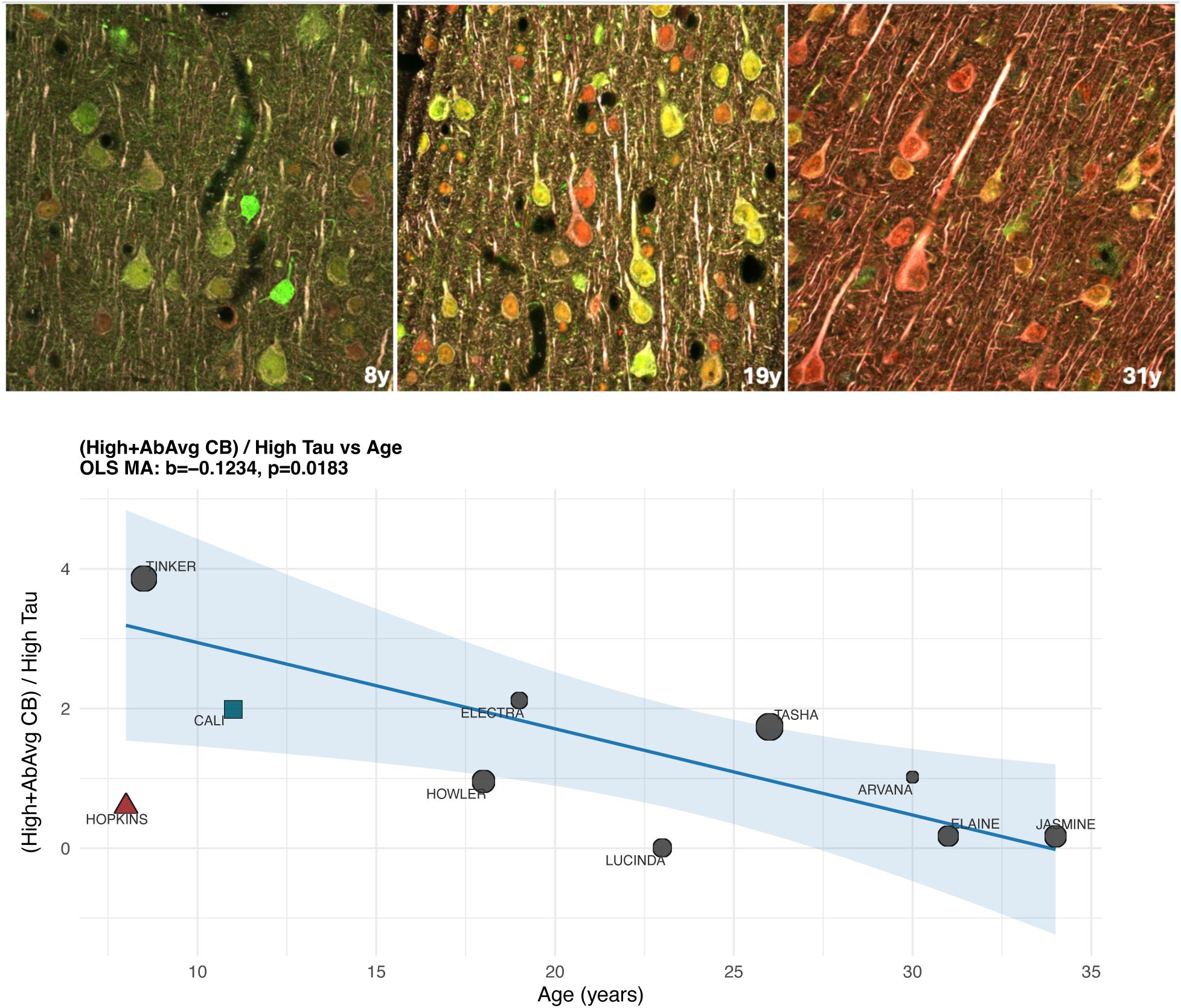
Panels A-C highlight representative merged images from a middle aged adult (A), early aged (B), and old aged (C), animal respectively. CB (green) is most prominent in the youngest of the animals, whereas pT217Tau (red) is most prominent in the oldest, both within the dendrites captured for analysis as well as throughout the neuropil. Structural marker MAP2 shown in white. The same pattern of positivity is captured by the analysis (D), which highlight that elevated CB to high pT217-tau ratio decreases across age span. Scale bars = 40um.

## DISCUSSION

This study provides quantitative evidence linking the age-related loss of calbindin-D28K with the accumulation of pT217Tau within individual dendritic compartments of dlPFC layer III pyramidal cells across the adult rhesus macaque lifespan. Using two complementary metrics applied to the same dendritic compartment, per-dendrite fluorescence intensity and area-occupied analysis, a progressive, age-related shift from high-CB to high-pT217Tau occupancy was observed, with all three CB/pT217Tau area-occupied ratios declining significantly following site correction (FDR-p=0.046). The loss of CB and rise in pT217Tau was particularly evident in the aged macaques with chronic inflammatory conditions. These findings have direct implications for understanding the molecular basis of early-stage tau pathology in circuits selectively vulnerable in AD. Since plasma pT217Tau is among the earliest detectable tau species in presymptomatic AD, reliably predicting future cognitive decline years before symptom onset^27^, the dendritic accumulation quantified here represents a likely subcellular contribution to this signal.

### Limitations

Several limitations warrant consideration. First, the cohort size (n = 10) limits statistical power for individual marker analyses and these data would benefit from replication in larger cohorts. The scarcity of aged rhesus macaques, exacerbated by demands of COVID-19 vaccine development, substantially constrains sample sizes in this model. Second, the cross-sectional design does not permit within-animal longitudinal tracking of CB and pT217Tau dynamics. Third, tissue was acquired at three sites, necessitating site correction; the youngest animal (Site 1) showed likely perfusion-related processing differences that attenuated the age effect when unadjusted, highlighting the critical importance of multi-site correction. Fourth, no systematically comparable cognitive assessments were available for this cohort.

### pT217Tau Accumulation in Aging Pyramidal Cells

The pT217Tau species quantified in this study accumulates at a biologically significant subcellular site and stage. Nanoscale immunoelectron microscopy in the aging macaque cortex has shown that soluble pT217Tau aggregates on microtubules within dendrites, where it might interfere with endosomal trafficking and is associated with autophagosomes, i.e. autophagic degeneration^9^. Critically, pT217Tau also appears to “seed” pathology between neurons at excitatory glutamatergic axospinous synapses, potentially exposing soluble tau to the extracellular space for capture in CSF and plasma^9^. It is important to note that tau phosphorylation is not inherently pathological. Tau normally functions to stabilize microtubules in dendrites and axons, and phosphorylation may allow dynamic structural plasticity, with subsequent dephosphorylation then allowing restabilization,^28,29^ consistent with the elevated plasma pT217Tau detected physiologically in infants^30^. With advancing age and inflammation, however, sustained calcium-activated kinase activity hyperphosphorylates tau beyond reversibility: rather than facilitating plasticity, accumulating pT217Tau destabilizes microtubules, impairs dendritic integrity, and initiates the pathological cascade toward neurofibrillary tangles^9^.

### Calcium Dysregulation Drives Tau Hyperphosphorylation

Extensive data show that aging, inflammation and Aβ_42_ all drive calcium dysregulation, which in turn drives tau hyperphosphorylation via activation of calpain-2^3,4^. Very high cytosolic calcium activates calpain-2, which cleaves and disinhibits GSK3β and p25-cdk5, two of the major kinases responsible for tau hyperphosphorylation^4,5^. Calpain-2 activation is an early, pervasive event in AD brains^31,32^, and its upregulation precedes tau pathology^33^. CB functions as one critical calcium buffer whose progressive loss can tip the balance toward elevated cytosolic calcium and resultant sustained calpain activation^13^. It should be emphasized that many other factors contribute to the rise in tau hyperphosphorylation; we do not imply that calbindin is the definitive factor. Nonetheless, it appears to be a key protective element that erodes with age and is especially reduced in animals with long-standing inflammatory disorders.

### Loss of Calbindin with Age, Inflammation, and in AD

The significant age-related decline in dendritic CB intensity observed in this study is consistent with prior evidence of progressive CB loss cells, in aging macaques^14^, and with age-related reductions in CB mRNA in human dlPFC^34^. Critically, CB is also vulnerable to inflammatory insults: patients who died from COVID-19 showed marked reductions in CB throughout the brain, including cerebellar Purkinje cells, accompanied by increased tau hyperphosphorylation^15^. In AD itself, the layer III dlPFC pyramidal cells that express CB in the healthy brain, are among the most vulnerable to tau pathology and degeneration in disease, while CB-retaining interneurons are relatively spared^17^. Early postmortem studies similarly reported loss of CB-immunoreactive neurons from the cortex as a feature of Alzheimer’s-type dementia^35^. Together, these convergent findings across species, aging trajectories, and inflammatory contexts implicate CB loss as a mechanistically relevant contributor to dendritic vulnerability.

### Individual Heterogeneity and Inflammation

These data are notable for the degree of individual heterogeneity observed, particularly in pT217Tau levels and CB/pT217Tau ratios. Although the sample size is quite small, our findings suggest that two animals with documented chronic inflammatory disorders, one with inflammatory bowel disease and renal disease (23yrs) and the other with type II diabetes (31yrs), showed the most pronounced CB depletion and pT217Tau accumulation, consistent with a model in which inflammatory signaling might accelerate calcium dysregulation beyond the trajectory of normal aging^5^. Conversely, the oldest animal (34yrs) showed relatively low levels of both CB and pT217Tau, suggesting that the age-related trajectory is not uniformly progressive and that individual variation in cumulative inflammatory burden, genetic factors, and environment may substantially modify dendritic vulnerability. This animal also had excellent cognitive abilities and was performing well within a week of illness and sacrifice, suggesting the possible existence of protective factors that can maintain the CB-tau balance independent of age. It is also possible that some neurons may be able to increase their production of CB to try to overcome early-stage insults from inflammation, oxidative stress, Aβ42 accumulation, and tau alterations^5,36^ at least at stages when neurons are still resilient; there was some evidence of this in the earlier-aged animals, where some dendritic compartments showed elevated levels of both calbindin and pT217Tau simultaneously. This heterogeneity mirrors the well-established individual variation in cognitive decline and neuropathological burden seen in aging humans and non-human primates^25,37^, and suggests that the CB/pT217Tau balance may capture an axis of vulnerability modulated by inflammation rather than solely by age.

It should also be noted that across the representative immunofluorescence images, there is a general qualitative visual decrease in CB and increase in pT217Tau signal with advancing age. This pattern likely reflects genuine biological signal, as well as some contribution from background labeling; immunoelectron microscopy has previously established that fine dendrites and dendritic spines are major subcellular compartments for pT217Tau and calcium-related protein expression in macaque cortex^9,14^, but at the confocal microscopic level these structures cannot be distinguished from background and were therefore not analyzed, representing a technical limitation that likely leads to underestimation of the true magnitude of the dendritic CB-tau imbalance.

### Calbindin Expression and Selective Neuronal Vulnerability

The pattern of CB expression across the brain provides important context for these findings and helps to explain selective neuronal vulnerability in AD. Neuronal populations that maintain high CB expression, including cortical and hippocampal interneurons, dentate gyrus granule cells, and cerebellar Purkinje cells, show remarkable resistance to tau pathology even in advanced AD^38^. Consistent with this protective role, an aged macaque in our prior study who retained very high levels of CB had no evidence of tau pathology^14^. Conversely, patients who died from COVID-19 lost CB from cerebellar Purkinje cells, a population that normally retains very high CB expression and is spared even in advanced AD, and simultaneously developed tau hyperphosphorylation in this region^15^. The layer II cell islands of the entorhinal cortex, which are among the earliest sites of calcium dysregulation^39^ and tau pathology in both AD and aging macaques, never express CB in either humans or macaques^40^, suggesting that the absence of this calcium-buffering protection from early adulthood may help explain their exceptional vulnerability. In contrast, the dlPFC layer III pyramidal cells studied here express CB in the healthy, younger brain, reflecting the high calcium demands imposed by the recurrent excitatory connectivity underlying working memory^23^, but progressively lose this protection with age and inflammation. This trajectory suggests that progressive CB loss converts a once protected population into one increasingly susceptible to pT217Tau accumulation, consistent with the Braak stage progression of tau pathology from entorhinal to association cortices^1^.

### Relevance to Treatment

Given the destructive effects of excessive cytosolic calcium, restoring calcium regulation may be a helpful strategy to reduce the toxic effects of inflammation and reduce the risk of sporadic AD^4,41,42^. A critical and unresolved question is why and how inflammation causes the loss of CB; developing strategies to retain CB in the aging brain represents a major therapeutic priority, as the dramatic depletion observed in response to inflammatory insults such as severe COVID-19 infection^15^ indicates that inflammation can substantially reduce CB expression. One well-characterized inflammatory mechanism of calcium dysregulation involves GCPII-mediated erosion of mGluR3 regulation: GCPII activity in aged macaque dlPFC strongly correlates with pT217Tau levels^12^, and chronic inhibition of GCPII in aged macaques significantly reduces pT217Tau in dlPFC, entorhinal cortex, and plasma, with no evidence of adverse effects^12^. These findings support the broader principle that restoring calcium regulation, whether by targeting inflammatory signaling upstream^43^ or through direct or indirect calpain-2 inhibition^44^, may be an effective strategy for protecting the aging brain from early-stage tau pathology.

## Acknowledgements

This work was completed as a portion of a dissertation submitted to fulfill in part the requirements for the Degree of Doctor of Philosophy at Yale University.

This work was funded by PHS grants R01AG061190 to AFTA; 1R21AG079145-01, 1R21AG102968-01, KL2 TR001862, Alzheimer’s Association Research Grant AARGD-23-1150568, Alzheimer’s Association and National Alzheimer’s Coordinating Center grant NIAP25-1445690, and P30AG066508 Developmental Project Award to DD; and T32 NS041228, T32 NS007224, and funding from the Gruber Science Fellowship to IP.

## Notes

### Competing Interest Statement

The authors have declared no competing interest.

