## Supplemental Methods for "Loss of calbindin and rise in pT217-tau in aging monkey prefrontal cortical dendrites"

**Supplemental Table 1.** Animals, ages, acquisition sites, and known health conditions.

| **Age (years)** | **Acquisition Site** | **Known Health Confounds** |
| --- | --- | --- |
| 8.02 | Site 1 | none |
| 8.06 | Site 3 | none |
| 11.28 | Site 2 | none |
| 18.35 | Site 3 | arthritis |
| 18.95 | Site 3 | none |
| 23.94 | Site 3 | Jejunal adenocarcinoma, chronic renal disease, inflammatory bowel disease, and endometriosis |
| 25.95 | Site 3 | Cardiac fibrosis, acute sepsis |
| 30.26 | Site 3 | Diabetes mellitus (controlled), vision loss |
| 30.80 | Site 3 | Diabetes mellitus (uncontrolled), endometriosis, severe weight loss |
| 34.49 | Site 3 | Arthritic, endometriosis |

### **S1. Multi-Site Correction for Area-Occupied Analysis**

Tissue for this study was acquired at three sites. To account for systematic immunofluorescence differences attributable to tissue processing across sites, acquisition site was modeled as a fixed additive covariate in all area-occupied statistical analyses. Specifically, OLS regression models took the form outcome ~ age + site, with Site 3 (n = 8 animals; ages 8.06–34.49 years) as the reference level. Site 1 (n = 1 animal; 8.02 years) and Site 2 (n = 1 animal; 11.28 years) were each included as binary indicator variables.

Because each singleton site contains a single animal, the site fixed effect for Sites 1 and 2 absorbs both acquisition-protocol differences and biological individuality simultaneously; these two sources of variation cannot be fully disentangled. Accordingly, site offsets for the animals acquired from Site 1 and Site 2 are reported as combined technical-plus-biological deviations from the Site 3 reference cohort rather than as pure technical corrections.

Site correction was of particular importance for CB/pT217-tau ratio outcomes. The 8 yr old from Site 1 showed substantially lower CB/pT217-tau ratios relative to Site 3 animals of comparable age (High CB / High pT217-tau offset: β = –0.770, SE = 0.284, p = 0.035; (High + Above Average CB) / High pT217-tau offset: β = –2.626, SE = 1.031, p = 0.044), consistent with a likely perfusion-related processing difference that attenuated high-intensity CB signal at this site. Before site correction, ratio p-values ranged from 0.08 to 0.18 (unadjusted); after site correction, all three CB/pT217-tau ratios declined significantly with age (p = 0.013–0.037; FDR-p = 0.037). The 11.28 yr old animal from Site 2 showed no significant site offset for any outcome (all p > 0.42), indicating that its measurements were broadly consistent with the Site 3 reference cohort.

Per-animal exact ages were computed from recorded birth and perfusion dates using the formula: age (years) = days between birth and perfusion / 365.25. All statistical analyses were implemented in R (version 4.x) using base lm() for OLS models.
